# Mechanism of Microbicidal Action of E-101 Solution, a Myeloperoxidase-Mediated Antimicrobial, and its Oxidative Products

**DOI:** 10.1101/394114

**Authors:** G. A. Denys, Neil C. Devoe, P. Gudis, M. May, R.C. Allen, J. T. Stephens

## Abstract

E-101 Solution is a first in class myeloperoxidase-mediated antimicrobial developed for topical application. It is composed of porcine myeloperoxidase (pMPO), glucose oxidase (GO), glucose, sodium chloride, and specific amino acids in an aqueous vehicle. Once activated, the reactive species hydrogen peroxide (H_2_O_2_), hypochlorous acid and singlet oxygen are generated. We evaluated the treatment effects of E-101 solution and its oxidative products on ultrastucture changes and microbicidal activity against methicillin-resistant *Staphylococcus aureus* (MRSA) and *Escherichia coli*. Time kill and transmission electron microscopy studies were performed using formulations with pMPO or GO omitted. The glutathione membrane protection assay was used to study the neutralization of reactive oxygen species. The potency of E-101 solution was also measured in the presence of serum and whole blood by MIC and MBC determinations. E-101 solution demonstrated rapid bactericidal activity and ultracellular changes in MRSA and *E. coli* cells. When pMPO was omitted, high levels of H_2_O_2_ generated from GO and glucose demonstrated slow microbicidal activity with minimal cellular damage. When GO was omitted from the formulation no antimicrobial activity or cellular damage was observed. Protection from exposure to E-101 solution reactive oxygen species in the glutathione protection assay was competitive and temporary. E-101 solution maintained its antimicrobial activity in the presence of inhibitory substances such as serum and whole blood. E-101 solution is a potent myeloperoxidase enzyme system with multiple oxidative mechanisms of action. Our findings suggest the primary site that E-101solution exerts microbicidal action is the cell membrane by inactivation of essential cell membrane components.

---

E-101 solution is a first-in-class topical myeloperoxidase-mediated formulation developed as an antimicrobial open wound wash solution. It is composed of two enzymes, glucose oxidase (GO) and porcine myeloperoxidase (pMPO) in an aqueous vehicle. Upon topical application of E-101 solution containing glucose, hydrogen peroxide (H_2_O_2_) is produced *in-situ* by GO that drives pMPO-dependent oxidation of chloride to hypochourous acid (HOCl). Once generated, HOCl (or its conjugate base OCl^−^ (pKa = 7.5)) participates in a diffusion-controlled reaction with a second H_2_O_2_ molecule to yield singlet molecular oxygen (^1^O_2_), a metastable electronically excited reactant with a microsecond lifetime. Singlet oxygen is a potent electrophilic oxygenating agent capable of reacting with a broad spectrum of electron-rich compounds (Fig 1). The bactericidal activity of the MPO antimicrobial system is enhanced by the selective binding of MPO to the surface of target microorganisms (1-3). *In vivo* and *in vitro* studies have shown E-101 solution exert potent and broad-spectrum microbicidal action against Gram-positive and Gram-negative bacteria including multi-drug resistant pathogens (4-7).

**FIG 1.**
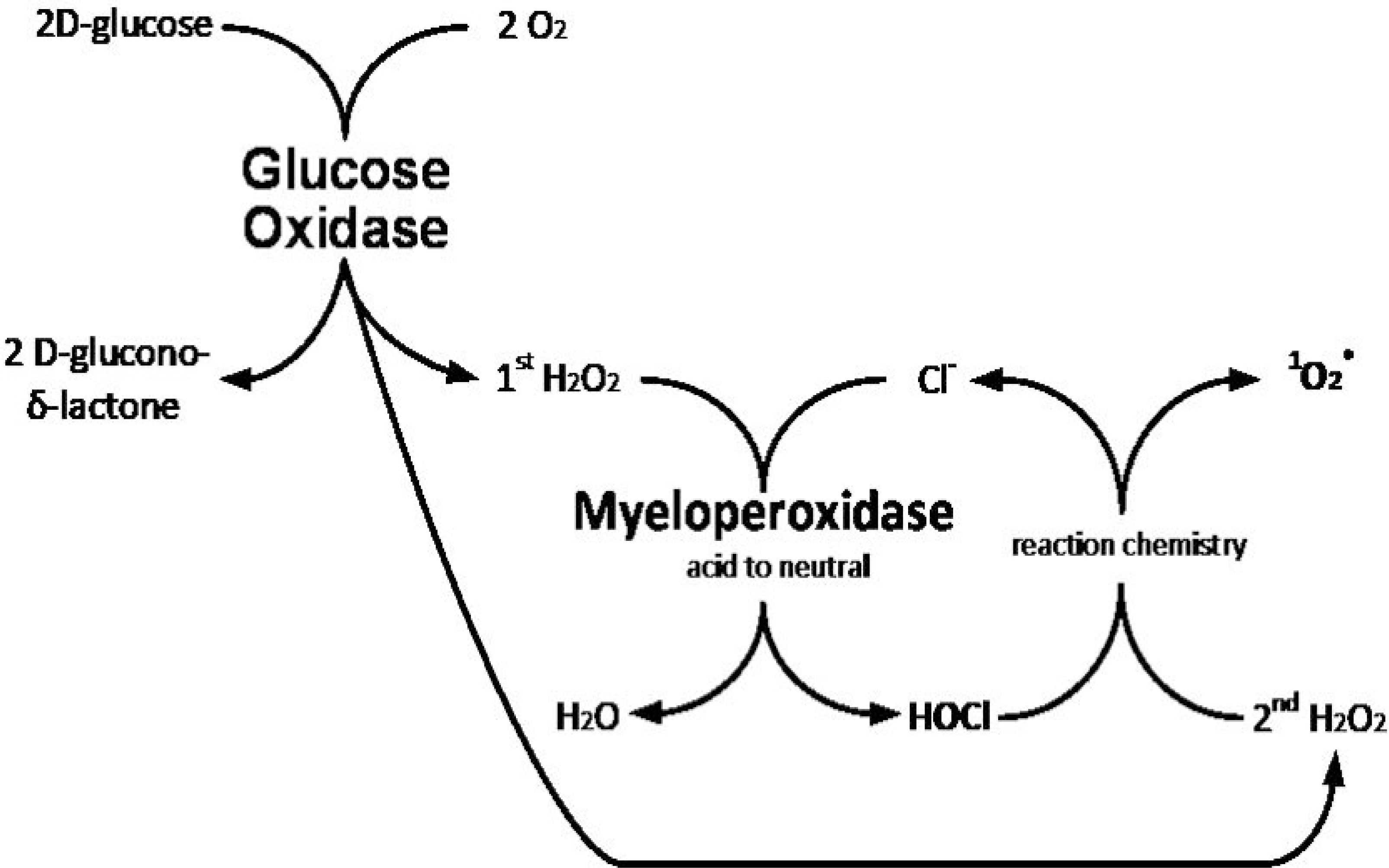
Schematic depicting the enzyme-linked oxidative action of E-101 solution. E-101 solution is composed of glucose oxidase (GO) and porcine myeloperoxidase (pMPO) in an aqueous vehicle and activated by the addition of glucose. H_2_O_2=_ hydrogen peroxide, Cl^−^ = chloride, HOCl = hyphochorous acid, ^1^O_2_ = singlet molecular oxygen.

The microbicidal combustive action of E-101 solution against target microorganisms is thought to be directed to a variety of molecular and enzymatic sites that are essential for metabolism or for the integrity of the microorganism (8). As part of new product development, this study was undertaken to better understand the potential mechanism(s) of action of E-101 solution and inhibition of activity. The goals of this study were: (i) to determine the time-kill effect of E-101 solution and its oxidative products on the ultrastructure of a Gram-positive and Gram-negative bacteria (ii) to determine the oxidative effect of E-101 solution on cellular damage using the glutathione membrane protection assay, and (iii) to compare the rate of kill of E-101 solution to sodium oxychlorosene in the absence and in the presence of serum and whole blood.

## RESULTS

### Time-kill effect of E-101 solution and oxidative intermediates

The bactericidal activity of E-101 solution was dependent on the entire components of the antimicrobial system (pMPO_−_+_−_H_2_O_2−_+_−_ halide). Time-kill curves of MRSA demonstrate rapid bactericidal activity of complete E-101 solution at 100, 416, and 833μg pMPO/ml (Table 1). At the early 5 min measurement, the rate of MRSA kill was inversely proportional to the E-101 solution concentration, but the 30 and 60 min measurements showed extensive 6-log kill. For *E. coli*, the 5 min and 30 min measurements of kill were directly proportional to the E-101 solution concentration, and at 60 min all concentrations showed extensive 6-log kill. The differences in early kill rates with respect to *S. aureus* and *E. coli* may reflect inhibition of MPO haloperoxidase activity by higher concentrations of H_2_O_2_ generated by GO at the higher E-101 concentrations tested. *S. aureus* has high catalase whereas *E. coli* has relatively low catalase expression. MPO is inhibited by concentrations of H_2_O_2_, especially if the ratio of H_2_O_2_ to chloride is high (9, 10)

**TABLE 1.** Comparative time kill of MRSA and *E. coli* exposed to complete E-101 solution, enzyme solution containing GO only, and enzyme solution containing pMPO only.

| Enzyme Solutions | Enzymes* |  | minutes | MRSA | E. coli |
| --- | --- | --- | --- | --- | --- |
|  | µg pMPO/ml | µg GO/ml |  | Log kill | Log kill |
| Complete E-101 | 100 | 20 | 5 | $1.5 \times 10^4$ | $1.2 \times 10^4$ |
| | 416 | 83 | 5 | $7.8 \times 10^1$ | $1.2 \times 10^4$ |
| | 832 | 167 | 5 | $1.3 \times 10^1$ | $2.3 \times 10^5$ |
| | 100 | 20 | 30 | $1.5 \times 10^6$ | $1.7 \times 10^4$ |
| | 416 | 83 | 30 | $6.7 \times 10^5$ | $1.2 \times 10^6$ |
| | 832 | 167 | 30 | $1.3 \times 10^6$ | $2.3 \times 10^6$ |
| | 100 | 20 | 60 | $1.5 \times 10^6$ | $1.2 \times 10^6$ |
| | 416 | 83 | 60 | $6.7 \times 10^5$ | $1.2 \times 10^6$ |
| | 832 | 167 | 60 | $1.3 \times 10^6$ | $2.3 \times 10^6$ |
| GO | 0 | 20 | 5 | 0 | <1 |
|  | 0 | 83 | 5 | <1 | <1 |
|  | 0 | 167 | 5 | <1 | <1 |
|  | 0 | 20 | 30 | 0 | <1 |
|  | 0 | 83 | 30 | <1 | <1 |
| | 0 | 167 | 30 | <1 | $2.7 \times 10^1$ |
| | 0 | 20 | 60 | <1 | $1.5 \times 10^3$ |
| | 0 | 83 | 60 | <1 | $2.0 \times 10^4$ |
| | 0 | 167 | 60 | <1 | $2.3 \times 10^6$ |
| pMPO | 100 | 0 | 5 | 0 | 0 |
|  | 416 | 0 | 5 | 0 | 0 |
|  | 832 | 0 | 5 | 0 | 0 |
|  | 100 | 0 | 30 | 0 | 0 |
|  | 416 | 0 | 30 | 0 | 0 |
|  | 832 | 0 | 30 | 0 | 0 |
|  | 100 | 0 | 60 | 0 | 0 |
|  | 416 | 0 | 60 | 0 | 0 |
|  | 832 | 0 | 60 | 0 | 0 |

\*Concentrations of pMPO tested were proportional to GO concentrations.

In the absence of pMPO, H_2_O_2_ generated from GO and glucose showed no activity against MRSA at 20μg GO/ml, but some antimicrobial activity at 83 and 167μg GO /ml, especially against *E. coli*. The bactericidal activity of H_2_O_2_ was observed at 167μg GO/ml, but at a slower rate (120 min). The pMPO-H_2_O_2_ microbicidal action was several orders of magnitude more rapid and more potent than H_2_O_2_ alone. The catalase content of *S. aureus* has been reported to be a virulence factor (11). *S. aureus* catalase competitively destroys H_2_O_2_ lowering its concentration without adversely affecting the rapid activity of E-101 solution. Sufficient H_2_O_2_ is present to drive the formation of both hypochlorous acid (HOCl) and singlet oxygen (^1^O_2_) which are thought to be the microbicidal agents in this system. However, in the absence of H_2_O_2_ generated by GO glucose catalyzed reduction of oxygen, pMPO is unable to exert any antimicrobial activity against catalase positive MRSA.

Time-kill curves of *E. coli* ATCC 25922 (Table 1) demonstrate similar rapid bactericidal activity of complete E-101 solution (pMPO + GO) at all three pMPO concentrations. High levels of H_2_O_2_ generated from GO and glucose in the absence of MPO showed bactericidal activity, but at a magnitudinally slower rate compared to the complete MPO-containing E-101 solution. Unlike *S. aureus, E. coli*, a weak catalase producer, showed susceptibility to H_2_O_2_ generation over time and at high concentrations. Formulation without GO added demonstrated no antimicrobial activity at all pMPO concentrations tested.

### Ultrastructure effect of E-101 solution and oxidative intermediates

Exposure of MRSA to complete E-101 antimicrobial solution at concentrations of 416 and 833μg pMPO/ml: 83 and 167 μg GO/ml for up to 120 minutes induced morphological changes (Fig 2). Both time-kill and TEM studies revealed that complete E-101 solution generates oxidative products that damage cells in a time- and concentration-dependant manner when pMPO, halide and a source of hydrogen peroxide are present. At 30 min, the majority of cells appeared ultrastructually normal, i.e., the cell wall, cell membrane and major constituents of the cell cytoplasms appeared similar to the control, but the microbes were killed as measured by CFU activity. However, post-kill ultrastructural changes were noted after 1 and 2 hours mainly at the cell membrane level. At 60 and 120 minutes, mesosome-like structures and septal defects in MRSA appeared. At 120 min, septal defects were more pronounce displaying thickened septa. Changes in the cytoplasmic membrane, mesosome-like structures, did not occur in untreated *S. aureus* cells. Increased vacuolation of the cytoplasm was also seen at 60 and 120 min. Similar morphological changes were observed in antibiotic treated *S. aureus* (12, 13). No apparent ultrastructure changes were observed in cultures treated with enzyme formulation when GO or pMPO were omitted.

**FIG 2.**
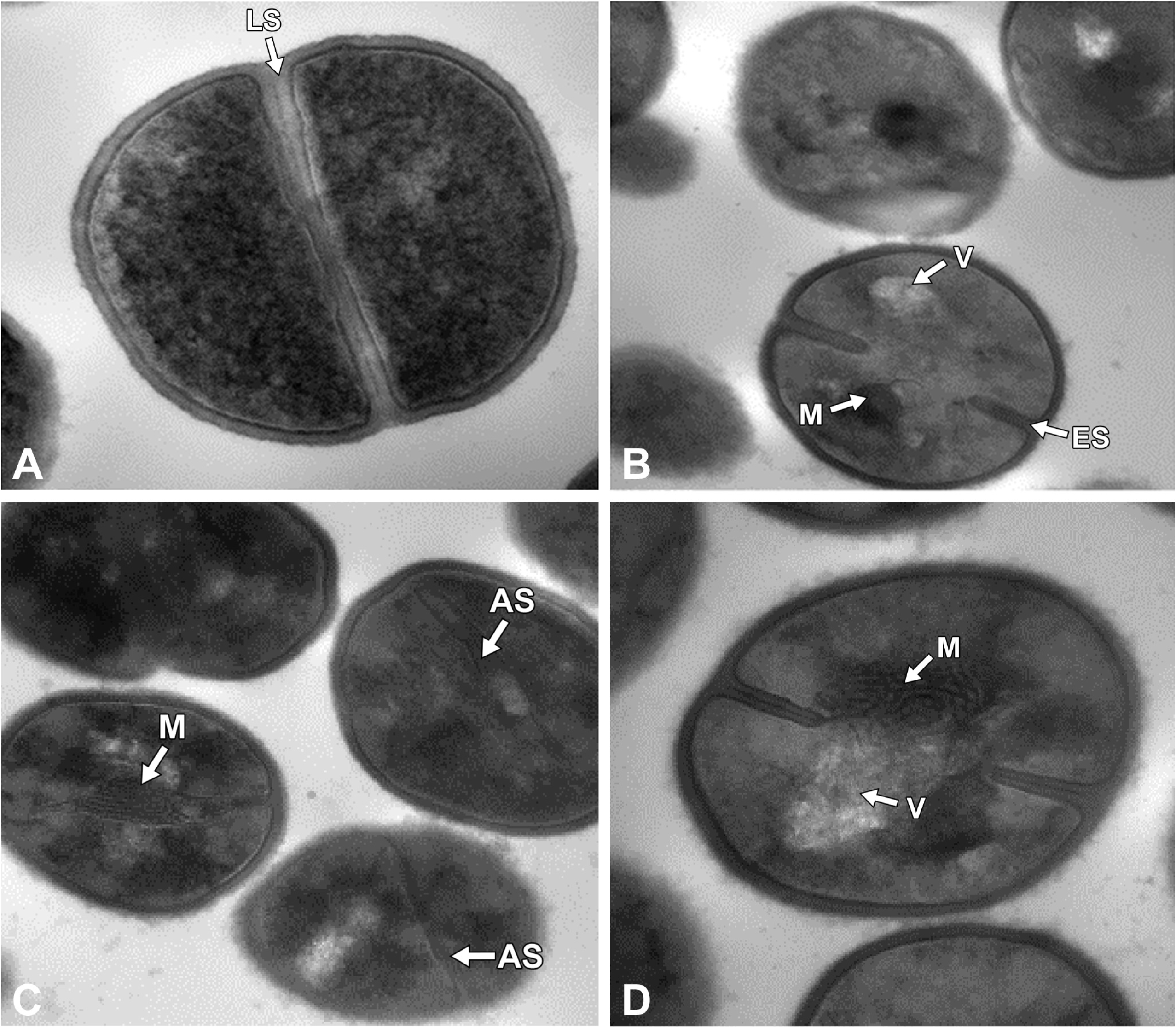
Morphological changes of MRSA cells harvested from exponential growth phase exposed to complete E-101 solution after 60 to 120 minutes. (A) Untreated control cells of MRSA showed uniform septa. Arrow indicates late uniform septa (LS), magnification: 68000x. (B) MRSA treated with complete E-101 solution at 120 minutes. Arrows indicate early septa (ES), cytoplasmic vacule (V), and cytoplasmic mesosome (M), magnification: 49000x. (C) MRSA treated with complete E-101 solution at 60 minutes. Arrows indicate aberrant septa (AS) and cytoplasmic mesosome (M), magnification: 49000x. (D) MRSA treated with complete E-101 solution at 120 minutes. Arrows indicate cytoplasmic vacule (V) and cytoplasmic mesosome (M), magnification: 68000x.

Exposure of *E. coli* to complete E-101 also induced time- and concentration-dependent morphological changes in septal formation and cytoplasm at 60 and 120 minutes (Fig 3). E-101 solution induced cellular elongation (septal deformation), cytoplasmic vacuoles, and pleated cell walls in *E. coli*. These post-kill findings are consistent with the activity of E-101 and the generation of reactive species H_2_O_2,_ HOCl, and ^1^O_2._ Cultures treated with partial formulation containing enzymes GO or pMPO alone demonstrated increase vacuolization. No abnormal cytoplasmic membrane or cell wall changes were observed when pMPO or GO was omitted from the formulation. The lack of obvious ultrastructure changes but extensive 6-log kill at 30 minutes of treatment with E-101 solution suggest that killing involves the combustive denaturation of key enzymatic components and/or destruction of membrane integrity that precedes ultrastructural damage.

**FIG 3.**
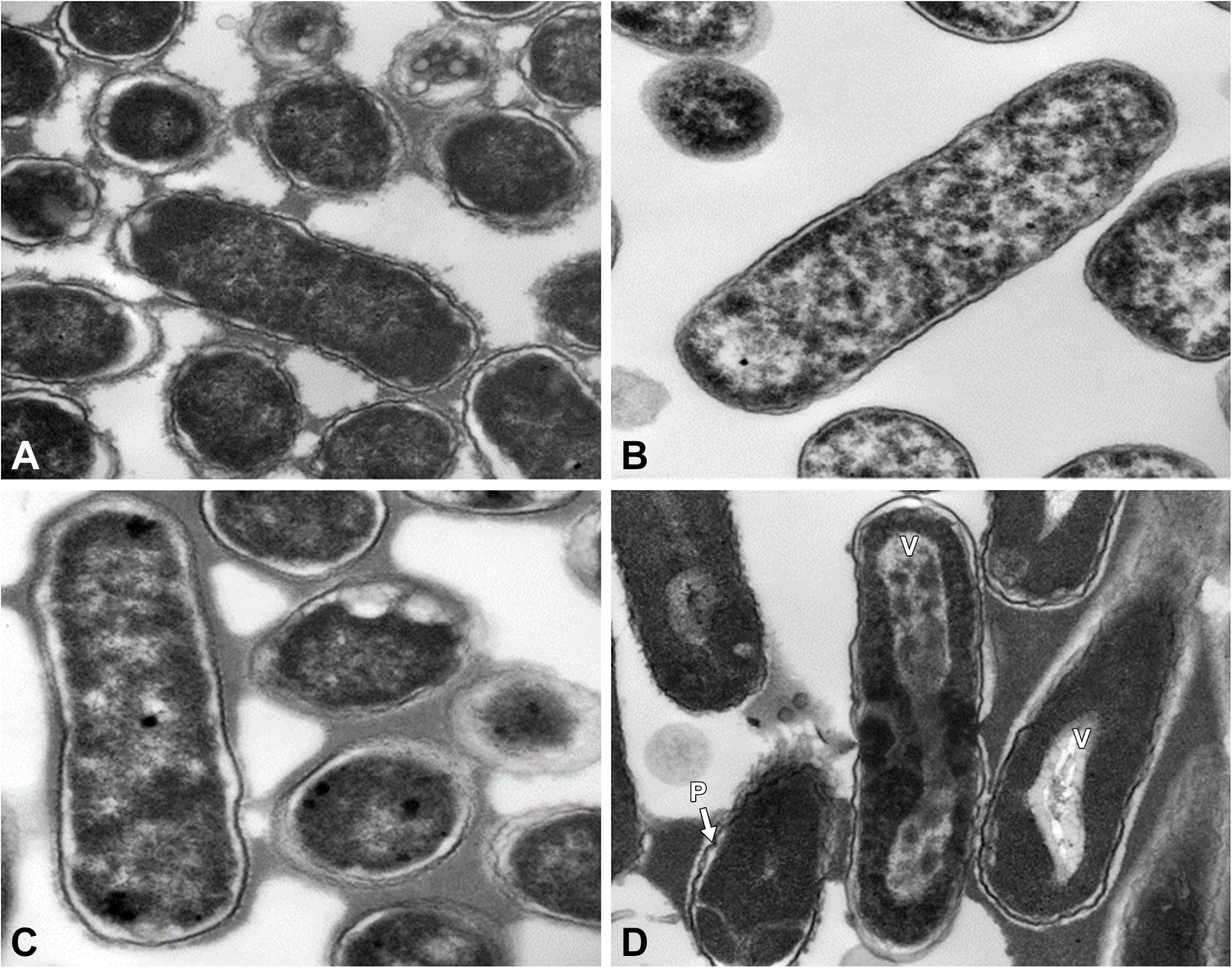
Morphological changes of *E. coli* cells harvested from exponential growth phase exposed to complete E-101 solution after 60-120 minutes. (E) *E. coli* untreated control at 0 minutes, magnification: 23000x. (F) *E. coli* treated with formulation containing GO (H_2_O_2_ generated) at 120 minutes, magnification: 30000x. (G) *E. coli* treated with formulation containing pMPO (no GO) at 120 minutes, magnification: 30000x. (H) *E. coli* treated with complete E-101 solution at 120 minutes. Arrows indicate pleated cell wall (P) and cytoplasmic vacule (V), magnification: 23000x.

### Oxidative destruction of glutathione by E-101 solution

The presence of 50 mM glutathione delayed but did not prevent E-101 solution killing of *S. aureus*. A significant (*P*<0.0001) reduction in viability was seen only after 10 minutes of exposure (FIG 4). The initial protective effect suggests that glutathione, a sulfhydryl reducing agent, neutralizes the oxidative and deoxygenating agents generated from E-101 solution The loss of inhibition after ten minutes suggests that once oxidative depletion of 50 mM glutathione is complete, oxidation of the target bacteria follows.

**FIG 4.**
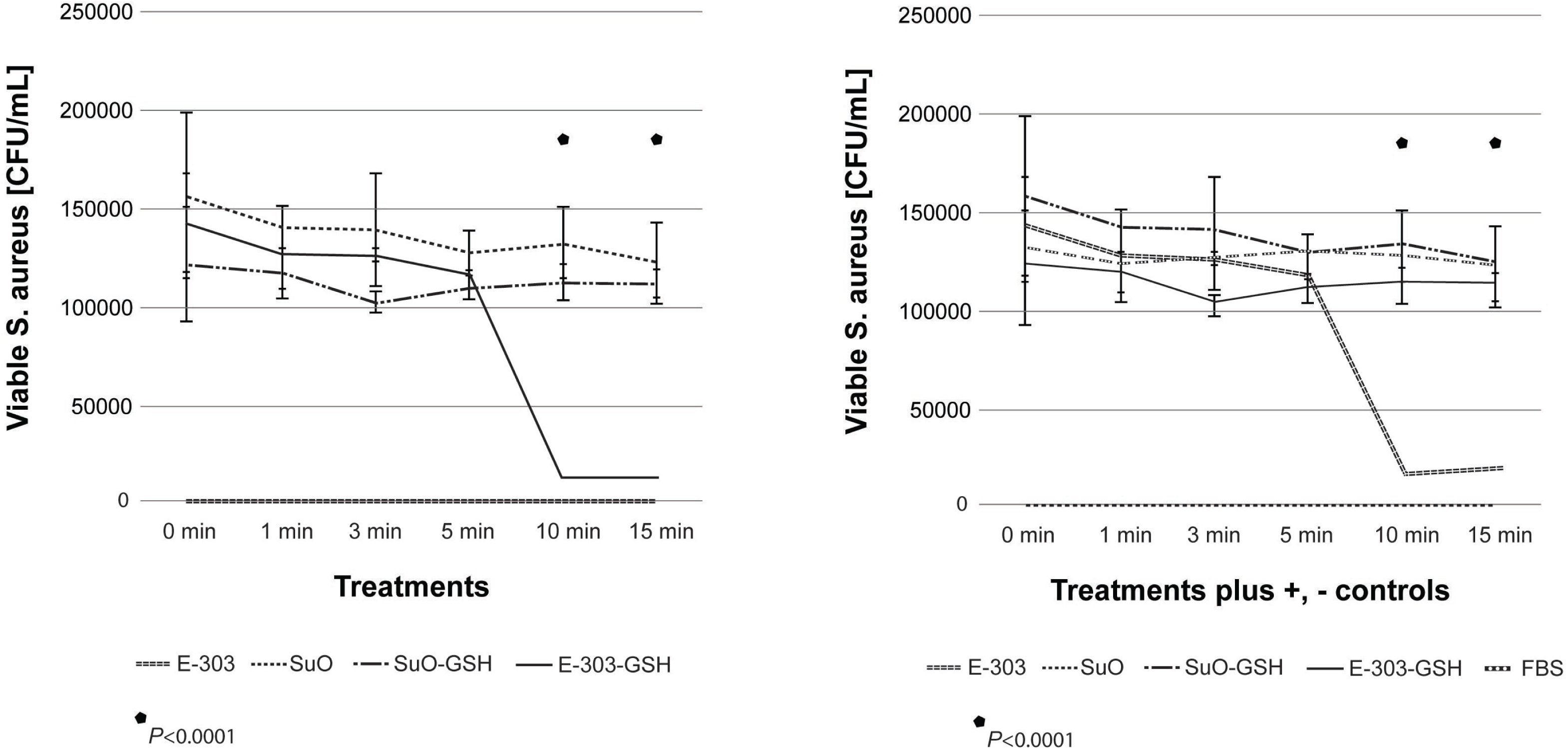
Glutathione membrane protection assay. (A, left) glutathione treatment. (B, right) glutathione treatment plus controls. Exposure of *S. aureus* to E-101 solution in the presence of 50 mM glutathione ablated the time required to achieve oxidative destruction of glutathione. A significant (*P*<0.0001) reduction in viability was seen only after 10 minutes of exposure compared to controls. E-101 = E-101 solution; Sub = E-101 substrate alone; Sub + GSH= E-101 substrate + glutathione; E-101 + GSH = E-101 solution + glutathione Positive controls: EtOH = ethyl alcohol; Negative control: PBS = phosphate buffered saline.

### Effect of inhibitors on myeloperoxidase antimicrobial system

The *in vitro* activity of the complete E-101 solution was compared to sodium oxychlorsene (Clorpactin), a buffered hypochlorous acid formulation, in the presence of serum and whole human blood. Table 2 summarizes the MICs and MBCs of both agents against MRSA and *E. coli*. Both E-101 solution and Clorpactin were highly active in the absence of serum or blood. However, in the presence of serum or blood E-101 solution maintained a high level of activity whereas Clorpactin activity was completely inhibited. The active component of Clorpactin is HOCl. Based on time-kill kinetic data in the absence of serum or blood, the rate of kill was slightly faster for Clorpactin (1 vs 15 min), however, E-101 solution demonstrated sustained and more potent kill than Clorpactin against *S. aureus* in the presence of 2, 5, and 10% blood (Fig 5). This is consistent with the selective binding and microbicidal action of MPO (1). The reaction of HOCl with an additional H_2_O_2_ yields ^1^O_2_ with a microsecond lifetime that restricts killing to the proximity of MPO binding. The potent oxidative anti-bacterial activities of E-101 solution was less susceptible than Clorpactin to the inhibitory effect of blood containing catalase and other competitive substances.

**TABLE 2.**
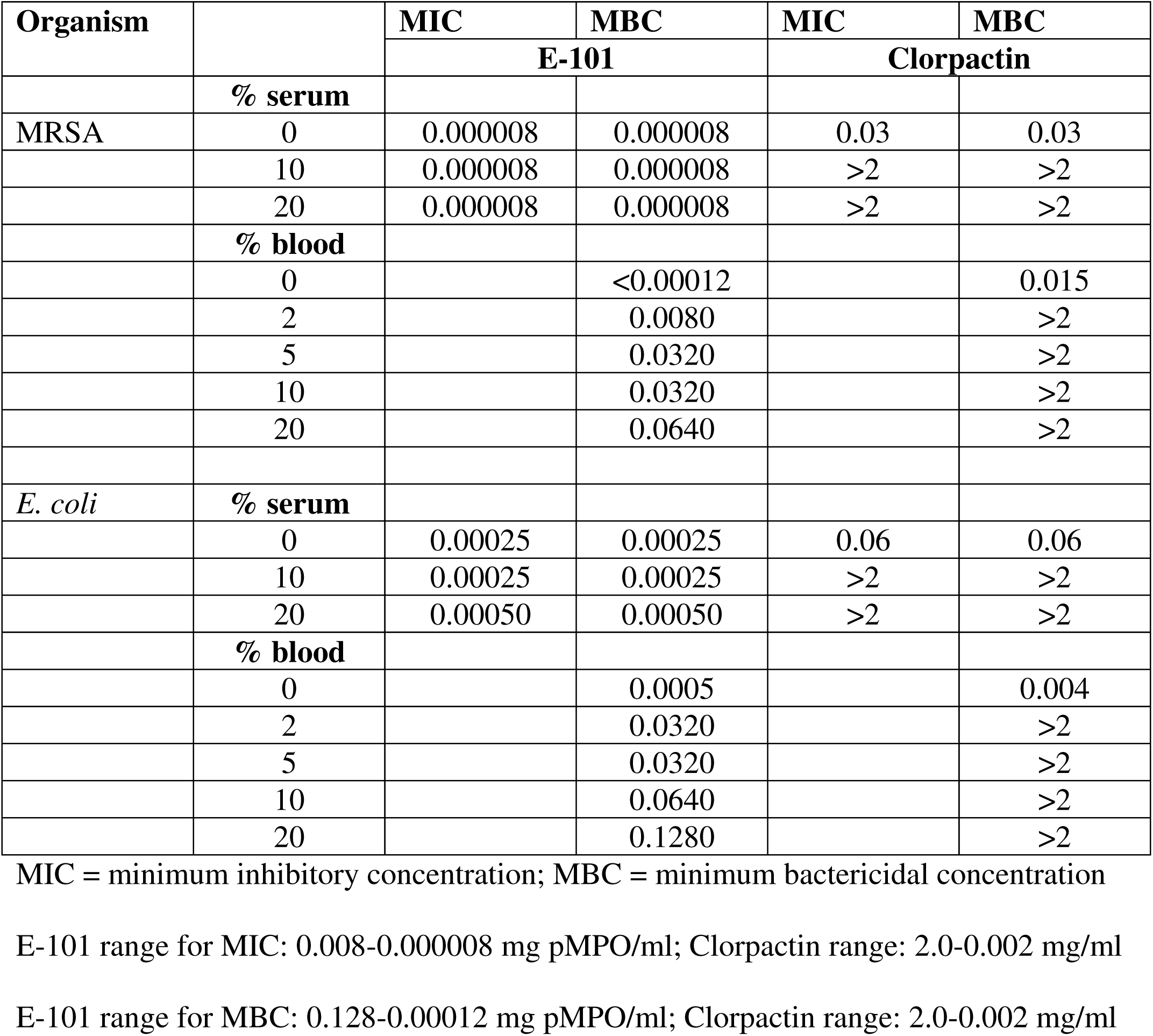
Comparative *in vitro* activity of complete E-101 solution and Clorpactin in the presence of horse serum and human whole blood.

**FIG 5.**
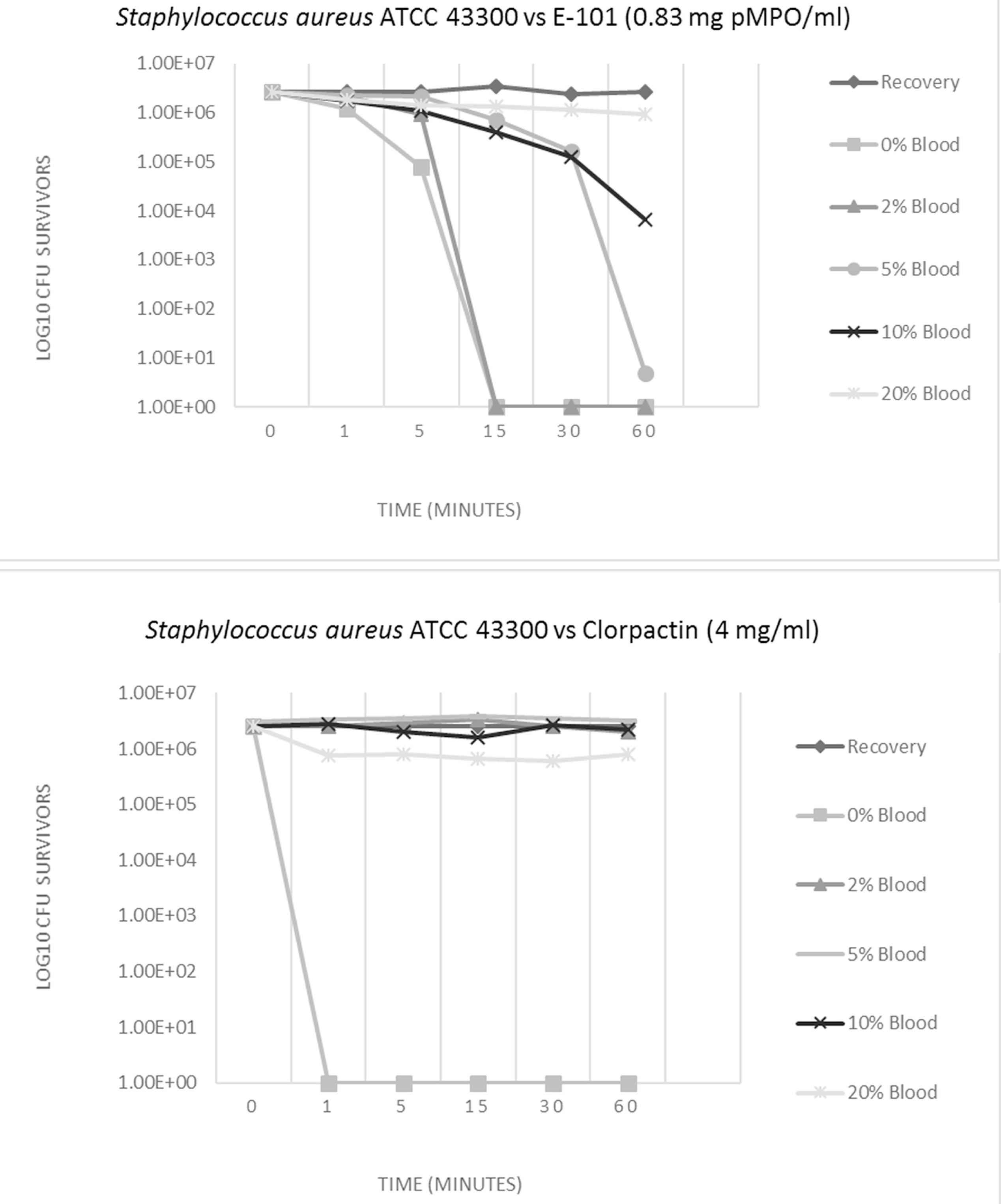
(top). Time-kill results for MRSA treated with complete E-101 at 0.83 mg pMPO/ml showed bactericidal activity (≥5-log_10_ reduction) in the presence of 2 and 5 % whole human blood with continued activity in the presence of 10% blood. No activity was observed in the presence of 20% whole human blood. FIG 5 (bottom). Time-kill results for MRSA treated with Clorpactin at 4 mg/ml showed rapid kill within one minute in the absence of whole human blood, but no activity in the presence 2% or greater whole human blood.

## DISCUSSION

Myeloperoxidase (MPO) is a heme-containing haloperoxidase present in high concentrations in neutrophilic granulocytes. MPO is one of the key components of the oxygen-dependent antimicrobial system of the neutrophil phagosome (14). The importance of MPO in killing of phagocytosed pathogens has been well documented (15, 16). The ability to efficiently kill phagocytized microbes requires the activities of membrane-bound nicotinamide adenine dinucleotide phosphate oxidase (NADPH oxidase) and myeloperoxidase (MPO) within the phagolysosome. The oxidase uses NADPH to reduce oxygen to ultimately generate the H_2_O_2_ that drives the MPO-dependent oxidation of chloride, yielding HOCl, and finally ^1^O_2_. Both HOCL and ^1^O_2_ are microbicidal agents in this antimicrobial system. (1, 17). Singlet oxygen formed by HOCl-mediated oxidation of H_2_O_2_ has been implicated as the principal bactericidal oxidant in the phagosome (18, 19). The highly electrophilic reactivity of ^1^O_2_ enables it to oxidize regions of high electron density in target biological molecules resulting in destruction of membrane integrity and/or the oxidative inhibition of the enzymes required for metabolic function.

*In vitro* studies confirmed that the combination of MPO, its substrate H_2_O_2_ and a halide form a potent antimicrobial system (20). The MPO-H_2_O_2_-halide system is capable of generating a wide range of oxidant species including ^1^O_2,_ OCl^−^, and chloramines (14). All of these products are reactive, but only ^1^O_2_ is electronically excited and thus short lived. OCl^−^ salts and chloramines can be stored for years. These highly reactive compounds can potentially oxidize susceptible reactive groups in biochemical substrates including thiols, thio-esters, heme groups, and unsaturated fatty acids (21). Such oxidative modifications alter protein and lipid membrane activities and consequently affect microbial and cellular functions including membrane structure and metabolic activity.

E-101 solution is a formulated cell-free oxidant-generating enzyme system, which mimics the intrinsic functions of the phagolysosome *in situ*. It consists of two enzymes; porcine-myeloperoxidase (pMPO) and glucose oxidase (GO), glucose, which is the substrate for GO, sodium chloride, and proprietary amino acids that enhance the activity of the system once it has been activated after mixing all of the components. In this study, the microbicidal activity of E-101 solution was dependent on each component of the functional system. The absence of pMPO resulted in magnitudinally diminished antimicrobial activity. The absence of GO also resulted in impaired antimicrobial activity. The microbicidal mechanism of action by E-101 solution involves binding of pMPO to the surface of target microorganisms where hypochlorous acid (HOCl) and singlet oxygen (^1^O_2_) focus direct oxidative damage. (1, 2, 4, 22, 23). In the presence of hydrogen peroxide (H_2_O_2_), generated *in-situ* by GO from glucose and oxygen, the microorganism bound pMPO catalyzes the oxidation of chloride ion to HOCl. Hypochlorite reacts with an additional H_2_O_2_ yielding ^1^O_2_ at or near the surface of the target organism. Although the glutathione membrane protection assay demonstrated initial protection from the reactive oxygen species generated from E-101 solution, complete killing of microorganisms after a lag time indicates that the protection provided by this reducing agent is competitive and temporary.

Ultrastructural evidence follows the microbicidal action. Time-kill results demonstrating that E-101 solution kills within minutes of direct exposure, and that this microbicidal action occurs prior to any observable disruption of the microbial plasma membrane suggesting oxidative inhibition of vital membrane enzyme systems. E-101 solution induced subtle, but clearly visible changes in bacterial cells at the longer incubation times. Active pMPO was essential in demonstrating cell damage since no damage was seen when pMPO was omitted from the formulation. Similar studies support these finding in damaging Candida hyphae and pseudohyphae. Oxidative intermediates appeared to damage hyphae only if MPO, halide and a source of H_2_O_2_ are present or if ^1^O_2_ is produced (24).

Consistent with the selective binding and focused combustive oxygenation mechanism of action, E-101 solution was less susceptible to the inhibitory effect of blood containing catalase and other substances that competitively react with available ^1^O_2_ and HOCl (10). Microorganisms with strong MPO binding capability are targeted for MPO-generated HOCl and ^1^O_2._ In contrast, Clorpactin with the active component HOCl was rapidly bactericidal but easily inactivated by the presence of serum or blood. Previous studies demonstrated that the microbicidal activity of MPO-H_2_O_2_ is several orders of magnitude more potent than that of H_2_O_2_ alone and more resistant to erythrocyte inhibition than the activity of either H_2_O_2_ or HOCl (1-2).

In summary, our data demonstrate that the bactericidal action of E-101 solution are attributed to the MPO-derived oxidants generated from the pMPO-H_2_O_2_-Cl antimicrobial system. E-101 solution reacts with the bacterial membrane components that are essential for viability and cell division. This enzyme system may play an important role in antimicrobial therapy.

## MATERIALS AND METHODS

### Reagents and enzymes

Stock solutions of complete E-101 solution containing pMPO and GO, Enzyme solution containing GO, Enzyme solution containing pMPO, and Substrate solution containing glucose were prepared at Exoxemis, Inc. (Omaha, NE). A range of concentrations of Enzyme solutions were prepared just prior to use. The Complete E-101 solution contains pMPO; GO derived from *Aspergillus niger,* and proprietary amino acids in an aqueous formulation vehicle consisting of 150 mM sodium chloride and 0.02% w/v polysorbate 80 in pH 6.5, 20 mM sodium, and phosphate buffer. Enzyme solutions containing individual components of GO or pMPO alone were prepared in the same aqueous formulation vehicle. The stock concentrations of pMPO and GO were 2.5 mg/ml and 0.5 mg/ml, respectively. The Substrate solution contains 300 mM glucose in the same aqueous formulation as the Enzyme solution. The Enzyme and Substrate solutions are packaged in two separate vials and mixed together to activate the system. Activated formulations were held at room temperature for 15-20 minutes for oxidant generation before use.

Sodium oxychlorosene (Clorpactin WCS-90) was obtained from Guardian Laboratories (Hauppauge, NY) and prepared according to manufacturer’s directions. Donor herd horse serum was obtained from Sigma-Aldrich, Inc. (Milwaukee, WI). Volunteer donor whole blood was obtained from the Blood Bank Department at Indiana University Health Pathology Laboratory (Indianapolis, IN).

### Time-kill assay

The microbicidal effect of complete E-101 Solution and Enzyme solutions against methicillin-resistant *Staphylococcus aureus* (MRSA), ATCC 43300, and *Escherichia coli,* ATCC 25922, were determined by a modified CLSI time-kill assay Several colonies (total 4 to 6), grown on TSA with 5% sheep blood overnight, were suspended in 3 ml of 0.45% sodium chloride and suspensions were adjusted to a 1.0 McFarland standard. This suspension was then diluted 1:10 in pre-warmed trypticase soy broth (TSB) and incubated for 2 to 4 hours at 35°C. When the culture reached its logarithmic growth phase and the turbidity approximated that of a 1.0 McFarland standard, the suspension was diluted 1:5 in normal saline (final concentration ∼ 6.0 × 10^7^ CFU/ml). This suspension constituted the inoculum for the time-kill assay. The *in vitro* assays were conducted in glass tubes (20 × 125 mm). The test-kill reaction tubes were prepared to contain 4.5 to 5 ml of appropriate inoculum in saline, substrate and enzymes (complete E-101, GO or MPO alone). Complete E-101 solution was mixed 15 minutes prior to its use. The final concentrations of pMPO in complete E-101 solution and pMPO solution alone were 100, 416, 833 μg/ml (36, 150, and 300 GU/ml). The final concentrations of GO in complete E-101 solution and GO alone were 20, 83, and 167 μg/ml. An identical reaction tube containing saline plus inoculum and Substrate solution, but no enzymes, constituted the culture growth control. The final bacterial cell concentration was ∼10^6^ CFU/ml. The reaction tubes were incubated at 35° C in ambient air and samples were removed for viable counts at 0, 5, 15, 30, 60, 90 and 120 minutes. Serial samples were obtained for quantification. A 100 μL-sample was immediately removed from each tube at each time point and applied to TSA with 5% sheep blood and serial 10-fold dilutions were prepared in sterile saline. A 100 μl-volume of each dilution was applied to duplicate TSA with 5% sheep blood plates and spread over the surface with a sterile inoculating loop. The plates at time zero functioned as the purity plates. Following overnight incubation at 35°C, colonies were manually counted and viable counts were calculated. The data are presented as log_10_ reduction in cfu/ml at designated time points compared to the original cfu/ml at the start of testing. Bactericidal activity was defined as a 99.9% or a 3-log10 cfu/ml reduction in the colony count from the initial inoculum.

### Transmission Electron Microscopy (TEM)

Several colonies (total 4 to 6) of MRSA, ATCC 43300, and *E. coli*, ATCC 25922, grown on TSA with 5% sheep blood overnight, were suspended in 3 ml of 0.45% sodium chloride and suspensions were adjusted to a 1.0 McFarland standard. This cell suspension was then diluted 1:10 in pre-warmed TSB and incubated for ∼2 to 4 hours at 35°C to logarithmic phase growth. The cell suspension was centrifuged (3,600 rpm for 6 minutes), washed with normal saline, and resuspended in normal saline to a 3 McFarland standard (∼10^9^ CFU/ml). Control cells consisted of equivalent volumes of organism suspension and normal saline with no test article added. Equivalent volumes of organism suspension (e.g. 3 ml) and enzyme formulation (e.g. 3 ml) were combined at the desired concentration (e.g. 100, 416, 833 μg/ml (36, 150 and 300 GU/ml). Treated cell suspensions were then incubated at 35° C and sampled at 0, 30, 60 and 120 minutes. Cells were harvested by centrifugation (10,000 rpm for 10 minutes) and fixed in 3.0% (v/v) glutaraldehyde.

Pellets were fixed for 24 hours (minimum) in 3% glutaraldehyde buffered with 0.15M Na cacodylate pH 7.2. Following 3 rinses in cacodylate buffer the pellets were post fixed in 1% osmium tetroxide (aq) for 4 hours at room temperature. The pellets were rinsed in buffer and dehydrated through a graded series of ethanol, rinsed twice in propylene oxide and left overnight in a 1/1 mixture of propylene oxide and Spurrs/Polybed epoxy resin. The pellets were infiltrated with full strength resin for 8 hours prior to embedding in beam capsules with fresh resin. The specimens were polymerized overnight in a 70^°^ C oven. Thin sections of the pellets were placed on 200 mesh cooper grids and stained with lead citrate and 3% uranyl acetate (aq). Sections were examined with a FEI Tecnai G^2^ Spirit transmission electron microscope operated at 80 kv. Images were captured with an AMT XR 60 digital camera.

### Glutathione membrane protection assay

Complete E-101 solution was mixed 15 minutes prior to its use. One ml of E-101 active solution was added to 2 ml substrate solution and left to acclimate to room temperature per manufacturer’s instructions. An inoculum of *S. aureus* strain, ATCC BAA 1717, was cultured for 4 hours to generate actively replicating bacterial cells within a standard incubator. 50 μl of the culture was added each 3 ml condition tube. Time kill assays were performed by exposing bacteria for 0, 1, 3, 5, 10 & 15 minutes for each of the following conditions: E-101 treatment; E-101 substrate alone; E-101 plus glutathione buffer (50 mM glutathione, 75 mM sodium chloride, 1 mM magnesium chloride, 0.1 mM calcium chloride, 80 mM sodium hydroxide, and 10% FBS, pH 8.6 (26); and E-101 substrate plus glutathione buffer. Treatment with 80% ethyl alcohol (EtOH) and phosphate buffered saline (PBS) served as positive and negative controls, respectively.

Following exposure of *S. aureus* to each condition for the designated time, the reaction was quenched with the addition of 50 μl of catalase. Bacteria surviving after treatment were enumerated by 10-fold serial dilutions in brain-heart infusion (BHI) broth followed by inoculation of BHI agar. Plates were incubated for 24 hours, after which viable colonies were counted to assess survival rate. Bacteria are reported as colony-forming units per mL (cfu/ml). This study was performed in triplicate for statistical analysis by analysis of variance.

### MIC determination

Comparative minimum inhibitory concentration (MIC) activity against MRSA, ATCC 43300, and *E. coli*, ATCC 25922, were determined using a modified broth microdilution method based on CLSI M7 guidelines (27, 5). Modifications included diluting enzyme solutions containing pMPO in double strength cation-adjusted Mueller-Hinton broth (CAMHB). Next, a standardized bacterial suspension was prepared in double strength glucose substrate solution containing serum or blood and mixed with serial enzyme dilutions in CAMHB to achieve a final concentration of 5 ×10^5^ CFU/ml. Similarly, Clorpactin was diluted in double-strength CAMBH, and mixed with inoculum containing serum or whole blood. The microdilution panels were incubated in ambient air at 35°C for 18-24 hours. MICs were determined by observing the lowest concentration of antimicrobial agent that inhibited growth of the organisms. Organisms were tested in the presence of increasing concentrations of horse serum (0, 10, and 20%) and human whole blood (0, 2, 5, 10, and 20%). The test range for complete E-101 solution was 0.128 to 0.00012 mg pMPO/ml and test range for Clorpactin was 2 to 0.002 mg/ml.

### MBC determination

Comparative minimum bactericidal concentration (MBC) activity against MRSA, ATCC 43300, and E. coli, ATCC 25922, were determined in the presence of horse serum and whole blood. A10 μl sample from the last well of the MIC panel with visible growth and each clear well was plated onto TSA with 5% sheep blood. After incubation in ambient air at 35°C for 18-24 hours, plates were examined for growth and colony counts. MBCs were defined as a 99.9% or a 3-log10 cfu/ml reduction in the colony count from the initial inoculum.

### Comparative time assay in the presence of serum and whole blood

The time kill kinetic studies were conducted against MRSA, ATCC 43300, as described by Tote et al (28). Reaction tubes were prepared to contain logarithmic phase growth (10^6^ CFU), antimicrobial agent, and blood. The final concentration of activated complete E-101 solution (15 minute mix time) and Clorpactin were 0.83 mg pMPO/ml and 4 mg/ml, respectively. At desired contact times (1, 5, 15, 30, and 60 min.) the test mixture was added to neutralizer solution. After a 5 min incubation, the neutralized mixture was sampled for quantitative culture and incubated at 35° C for 24 hours. The log_10_ CFU at each time point was determined and compared to growth controls. Due to in-test dilution, both antimicrobial agents were tested at 80% of final doses.

#### Statistical analysis

The glutathione membrane protection assay test was performed in triplicate for statistical analysis by analysis of variance (ANOVA) at each time point using GraphPad Prism version 5.04.

## ACKNOWLEDGMENTS

We express appreciation to Michael Goheen for providing technical support during the electron microscopy studies. This work was presented in part at the 21^st^ European Congress of Clinical Microbiology and Infectious Diseases and 27th International Congress of Chemotherapy, Milan, Italy, 2011 (Abstract 1453) and 115^th^ General Meeting of the American Society for Microbiology, New Orleans, LA, 2015 (Abstract 2336). The glutathione membrane protection assay was performed at the University of New England College and Osteopathic Medicine, Biddeford ME, USA. This study was supported by Exoxemis, Inc., Little Rock, AR, USA.

